# Spatial covariation between wild boars and other mammals in peri-urban landscapes: insights for management from southern France

**DOI:** 10.1101/2025.05.26.656147

**Authors:** Laurine Barachin, Jules Dezeure, Sonia Saïd, Raphaël Mathevet, Eric Baubet, Simon Chamaillé-Jammes

## Abstract

Global urbanization is rapidly increasing, transforming ecosystems and favoring generalist species that adapt to human-modified environments. The wild boar, a highly adaptable species, is expanding its range and increasingly observed in urban and peri-urban areas, where it can cause accidents, damage, and health concerns. To mitigate their presence, non-lethal control strategies such as vegetation clearing in peri-urban parks and wastelands are being implemented to reduce the number of suitable resting sites. However, the ecological consequences of such habitat modifications on other wildlife species remain poorly understood. This knowledge gap limits the development of effective urban wildlife management strategies that seeks to balance ecological, social and economic considerations without harming wild boar populations and urban ecosystems.

In this context, we used camera trapping to study the covariation between wild boars and other mammal species in a peri-urban area, with the aim of guiding targeted and effective management strategies. Here, we show a widespread presence of wild boars but no spatial or temporal covariation between the wild boars and the other species. In contrast, we found strong spatial structuring in mammal communities, primarily driven by small carnivores (mustelids, genets, cats, foxes) and a minimal influence of temporal variation, confirming that spatial factors, rather than seasonal changes, largely shaped species distributions. The study of the temporal variation across months indicated minimal seasonal structuring but revealed substantial within-site variability in month-to-month species abundance, with a general abundance gradient driven by common species like wild boar, cat, and fox.

In conclusion, our results show that the lack of spatial covariation between wild boar and other species prevents reliable predictions, supporting a targeted rather than general approach to vegetation management.

## Introduction

The pace of global urbanization is accelerating. Projections indicate that urban areas will expand by a factor of 2.5 by 2030 (Angel et al., 2005), with 68% of the world’s population expected to live in urban areas by 2050 (United Nations, 2019). Urbanization is one of the main aspects of global anthropization, leading to major alterations of the ecosystems. Indeed, urbanization reduces and fragments patches of natural vegetation, destroying some species’ habitats, modifying others’ and creating new ones (Adams, 2005). Some species take advantage of these human-modified spaces, as they could be predator-free and/or have high resource availability (Fischer et al., 2015). Such species, known as synanthropes, adjust to living in human-modified landscapes such as cities and urban fringes by modifying their behavior (Ditchkoff et al., 2006; Ritzel and Gallo, 2020). They often become more nocturnal (Gaynor et al., 2018; Johann et al., 2020) and change their diet (Egert-Berg et al., 2021; Sugden et al., 2021). Therefore, synanthropes are typically generalist species (Ducatez et al., 2018), including the red fox (*Vulpes vulpes*), racoon (*Procyon lotor*) or wild boar (*Sus scrofa*) (Ballari and Barrios-García, 2014; Bateman and Fleming, 2012; Risch et al., 2021).

The wild boar is one of the most widely distributed mammals on Earth. Native to Eurasia and North Africa, it is now present on all continents except Antarctica (Long, 2003; Massei and Genov, 2004). Indeed, wild boar populations are increasing and their range is expanding (Beguin et al., 2016; Carpio et al., 2021; Côté et al., 2004; Massei et al., 2015; Snow et al., 2017). They have been observed at the edge of many major European cities (Licoppe et al., 2013), including Rome (Amendolia et al., 2019), Berlin (Stillfried et al., 2017b), and Barcelona (Cahill et al., 2012). The great phenotypic and behavioral plasticity of wild boars (Castillo-Contreras et al., 2021; Podgórski et al., 2013) explain their extensive geographic range and the diverse ecosystems they live in, including urban environments (Caspi et al., 2022). Nevertheless, the presence of wild boars in urbanized areas has given rise to a multitude of concerns. Indeed, they have been identified as a potential source of traffic accidents and, although very rare, they can directly harm humans and pets (Mayer et al., 2023; Zuberogoitia et al., 2014). Furthermore, the potential for zoonotic transmission (Fernández-Aguilar et al., 2018; Meng et al., 2009) and other forms of disturbance - such as damaging private and public gardens, and consuming garbage (Licoppe et al., 2013) - represent additional concerns.

Authorities from cities with high wild boar abundance often therefore seek to mitigate incursions of wild boars into their urban and peri-urban areas. One potential solution under consideration is the culling of the population. However, urban areas are usually not suitable for hunting activities, and city dwellers tend to be opposed to these lethal methods (Massei et al., 2011). Consequently, non-lethal strategies and management actions are being explored. One of these strategies is the clearing of the brushwood in green spaces such as urban public parks and industrial wastelands. These areas offer wild boars places to rest undisturbed during the day, allowing them to exploit the city at night. Removing these potential resting sites could ‘push’ wild boars further away from the city, and thus limit incursions in the most human populated areas. Although this approach has not been proven to be effective, to the best of our knowledge, it is implemented in cities such as Barcelona, as part of comprehensive management plans (Castillo-Contreras et al., 2018), and is considered elsewhere such as in Montpellier, where we work. A key question that arises when discussing such a management approach is what the consequences of vegetation clearing might be for the other wildlife species inhabiting these peri-urban areas. Indeed, the targeted areas may also be critical habitats for other animal species, including species with specific conservation and protection statuses. This knowledge gap hinders effective decision-making, as it remains unclear how to manage these habitats to reduce wild boar presence without negatively impacting other species.

Our study aimed at filling this gap by improving our understanding of the spatial and temporal covariation between wild boars and other species in peri-urban environments. This will contribute to the development of targeted and ecologically sound management strategies. We did so by using one year of data from 48 camera-traps deployed in the urban and peri-urban areas of Montpellier, the seventh most populated French city, where wild boars, which are abundant in the surrounding countryside, have recently established within the city itself.

## Material and methods

### Study area

The study was conducted in Montpellier (Occitanie, South of France) and its 12 neighboring municipalities (Castelnau-le-Lez, Clapiers, Grabels, Juvignac, Lattes, Lavérune, Mauguio, Montferrier-sur-Lez, Saint-Aunès, Saint-Clément-de-Rivière, Saint-Jean-de-Vedas and Villeneuve-lès-Maguelone). This area encompasses an area of 255.7 square kilometers and has approximately 430,000 inhabitants, 300,000 of whom reside in Montpellier itself (Insee, 2024). Situated in the south of France, Montpellier - the administrative center and largest city of the Hérault department - lies near the Mediterranean coast to the south and the Cévennes mountains to the northeast. The area is situated within a Mediterranean climate zone (*Csa* Köppen classification, Peel et al., 2007). The Hérault department hosts a substantial wild boar population and ranks among the top five French departments for annual harvest numbers (Barboiron, 2025), serving as an indicator of their abundance. The number of wild boars culled during the hunting season in Hérault has nearly doubled over the past two decades, reaching more than 26,000 individuals during the 2023/2024 hunting season (Barboiron, 2025). This trend reflects a growing population and may account for wild boars increasingly encroaching upon urban areas such as Montpellier.

### Data collection

From July 2023 to October 2024, 48 camera-traps (*CTs*; models: StealthCam STC-G42NG and BolyGuard SG560X) were installed at 48 sites across the study area, with one CT installed at each site. These sites were selected based on evidence of recent wild boar presence, including indirect signs and reports from local officials and community members.

All the CTs were situated within an 8-kilometer radius of the center of Montpellier (Fig. 1). They were installed in a variety of habitats reflecting the diversity of landscapes in the study area. These habitats included wastelands characterized by tall grasses, shrubs, and small trees; urban parks; forest patches; peri-urban Mediterranean ‘garrigue’ scrubland; densely vegetated riversides; and wetlands dominated by common reeds. The CTs were placed on identified animal trails, at a height (30-150 cm) and in a direction, determined by the specific configuration of each site (e.g. slope, proximity of trees to the trail, vegetation density, etc.), allowing the detection of wild boars and any other animals of a similar size. The CTs were programmed to take a burst of three consecutive images with a one-second interval when triggered, with an inactive delay of 5 seconds between two consecutive bursts. In the event of a trigger occurring within the 10-second interval following the most recent image capture, likely due to the same individual or group of individuals, the CTs were configured to initiate another burst. This continued until no further triggering activity was observed. All images captured as a result of the same initial trigger (i.e. <10s apart) were considered to belong to a unique sequence.

**Fig. 1.**
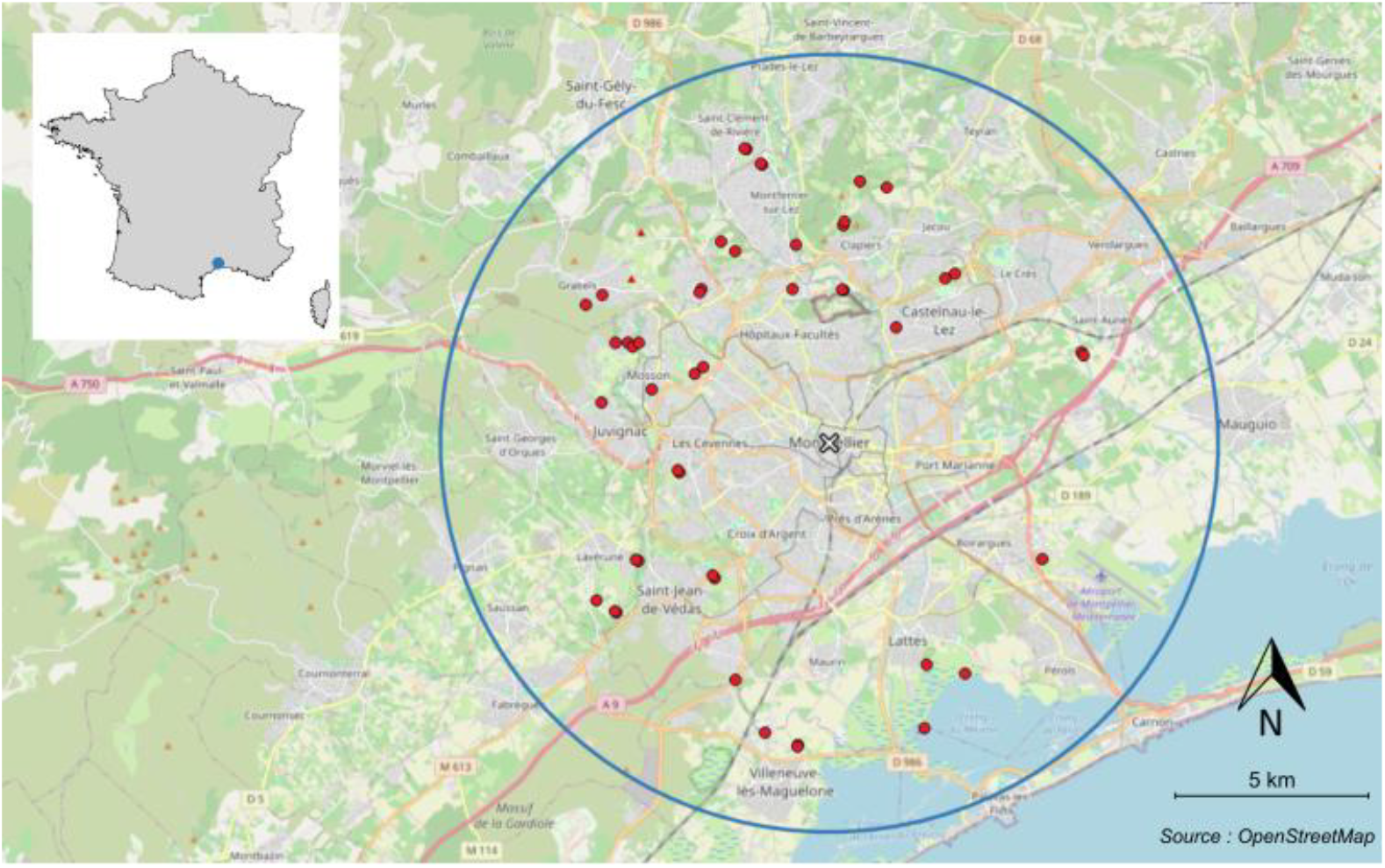
Location of the study area and the cameras traps. Location of the 48 CTs in the study area is shown in red diamonds and the blue concentric circle represents a radius of 10km from the center of Montpellier (white cross; 43°36’39”N 3°52’37”E).

### Species identification

We used artificial intelligence models to assist with species classification. The retrieved images were classified using the DeepFaune software (https://www.deepfaune.cnrs.fr; Rigoudy et al., 2023) but all classifications were visually checked and corrected when necessary. The number of animals present in each image was recorded. Each image sequence was classified into one of the following 25 classes: badger (*Meles meles*), cat (*Felis catus)*, goat (*Capra hircus*), roe deer (*Capreolus capreolus*), dog (*Canis lupus familiaris)*, squirrel (*Sciurus vulgaris*), equid (*Equus* species, for horses, poneys, donkeys), genet (*Genetta genetta*), hedgehog (*Erinaceus europaeus*), human (*Homo sapiens*), lagomorph (*Lagomorpha* order, for rabbits *Oryctolagus cuniculus* and hares *Lepus europaeus*), micromammal (brown rat *Rattus norvegicus*, wood mice *Apodemus sylvaticus*, vole species etc.), mustelid (*Martes* and *Mustela* genuses from the *Mustelidae* family, for pine marten *Martes martes*, stone marten *Martes foina*, least weasel *Mustela nivalis* etc.), bird (*Aves* class, including passerines, pigeons, corvids, raptors, picids and ardeids), butterfly (*Lepidoptera* order), coypu (also called nutria; *Myocastor coypus*), fox (*Vulpes vulpes*), wild boar (*Sus scrofa*), tortoise (*Testudo hermanni*), vehicle, empty (no animal), undefined (impossible to identify the animal), other (the observation belongs to none of the classes of DeepFaune), NA or error (linked to corrupted pictures). A total of 585,980 images were collected. After removing non-exploitable images (i.e., those classified as error, other, undefined, or NA), 581,034 images remained, corresponding to 164,586 independent sequences. As our analyses considered temporal variability, we focused our analyses on the period of consecutive weeks during which a minimum of 30 CTs were active (as sometimes less than 48 CTs were active due to non-synchronous deployments, battery failures, thefts…). Consequently, the dataset considered here was collected between the 1st of September 2023 and the 29th of August 2024, encompassing 52 weeks.

### Data exploration

As is commonly observed when using camera-traps, most (71.7%) of the image sequences were of the empty class. This high proportion was attributable to CTs being triggered by plant movements due to wind, light changes, strong winds moving the tree to which the CT was attached, and probably, sometimes animals such as birds passing quickly in front of the CT.

Classes that were not mammals (i.e. butterfly, bird, tortoise), or associated with human presence (human, vehicle, dog) were excluded from the dataset. Note that the cat class was retained, as cats move independently of people and as their important impact on biodiversity (Mori et al., 2019) is to be considered when discussing management options. Mammals with not enough observations (hedgehog, micromammal, squirrel, goat, equid) were also excluded. The animal classes retained for the study were therefore badgers, cats, roe deer, genets, lagomorphs, mustelids, foxes, coypus, and wild boars. This selection resulted in a dataset comprising 119,194 image sequences. For each of the nine classes, and at each site, we summed the number of individuals seen per day across all image sequences. Where the CTs were known to be non-functional, the number of animals was assigned the value “NA”. Subsequently, the data was summed per month for each CT, resulting in a tally of the number of individuals observed for each animal class per month and per CT. To account for the variability in the operation of the CTs, the number was weighted by the number of functioning days. For simplicity, we hereafter refer to this weighted monthly number only as “monthly number”.

### Statistical analyses

We studied the spatial and temporal covariations of wild boars with other mammals using Principal Component Analyses (*PCAs*) conducted on the monthly number of mammals seen at each CT. First, we performed a PCA on non-scaled centered data in order to quantify the absolute contribution of each species to the total variability observed in the data set. Secondly, we performed a PCA on centered and scaled data to examine their relative contribution to the observed variation. Both PCAs were performed using the *dudi.pca* function of the *ade4* package (Dray et al., 2023) of the R software (v.4.4.1; R Core Team, 2024).

To disentangle the relative importance of spatial versus temporal heterogeneity in the overall variability of the data, we carried out between- and within-group PCAs using the *bca* et the *withinpca* functions of the *ade4* package of the R software (Chamaillé-Jammes et al., 2016; Dolédec and Chessel, 1991). The spatial and temporal groupings were respectively based on the CT sites and the months. In each case, the total variability is decomposed in two additive parts, the between-group variation and the within-group variation. For example, with the spatial grouping, the total variability in species abundance observed within the dataset is decomposed into spatial variability (i.e. differences between CTs sites) and into non-spatial variability (i.e. differences within CTs sites).

Between-CT PCA identifies the species for which abundance varies the most between CTs, thereby emphasizing the differences between CTs in terms of spatial variation. Conversely, within-CT PCA eliminates the CT effect by centering the data by CT. This approach enables the identification of the primary structures that are not explained by the differences among CTs (i.e., non-spatial variation), which allows for the detection of temporal variability. The same line of reasoning can be employed with regard to between- and within-month PCAs. Thus, between-CT and between-month PCAs allow a direct estimation of the respective contributions of CTs and months to the overall variability. On the other hand, within-CT and within-month PCAs estimate the variations not explained by CTs and months respectively.

## Results

### Wild boar is the most seen mammal

Wild boars were detected at 92% of the sites (i.e. from 44 out of the 48 CTs). Of the 34,430 non-empty sequences obtained from our 48 CTs, wild boars were the most prevalent species, accounting for 60.0% of the sequences. This was followed by foxes and cats, with 15.5% and 14.9% of the sequences, respectively (Fig. 2).

**Fig. 2.**
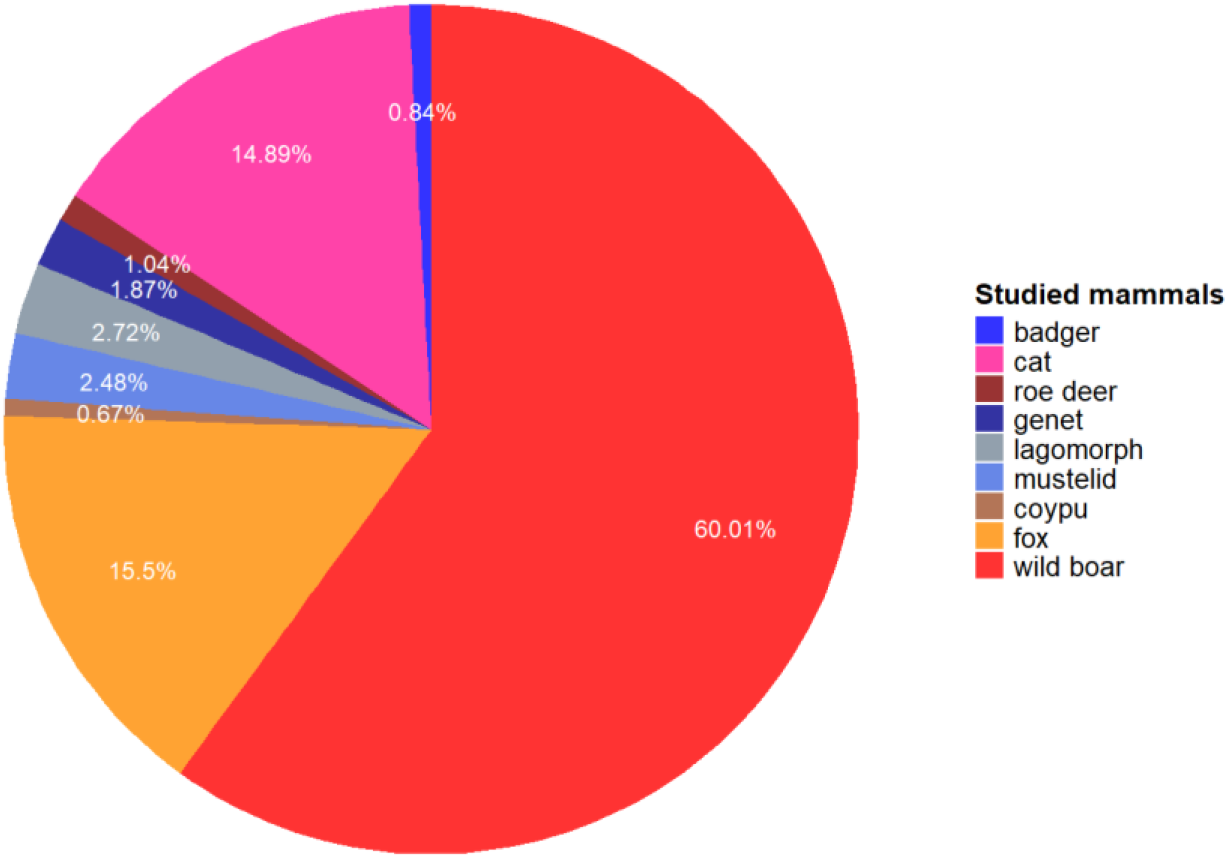
Proportion of camera-trap image sequences for the studied mammals.

Unsurprisingly given the above results, the first two axes of the PCA on non-scaled centered data captured the vast majority of the total variability (99%), with axis 1 alone accounting for 97.4% (Fig. 3) and being mostly associated with wild boar relative abundance. The second axis was mostly associated with the relative abundance of cats, and, to a lesser extent, foxes.

**Fig. 3.**
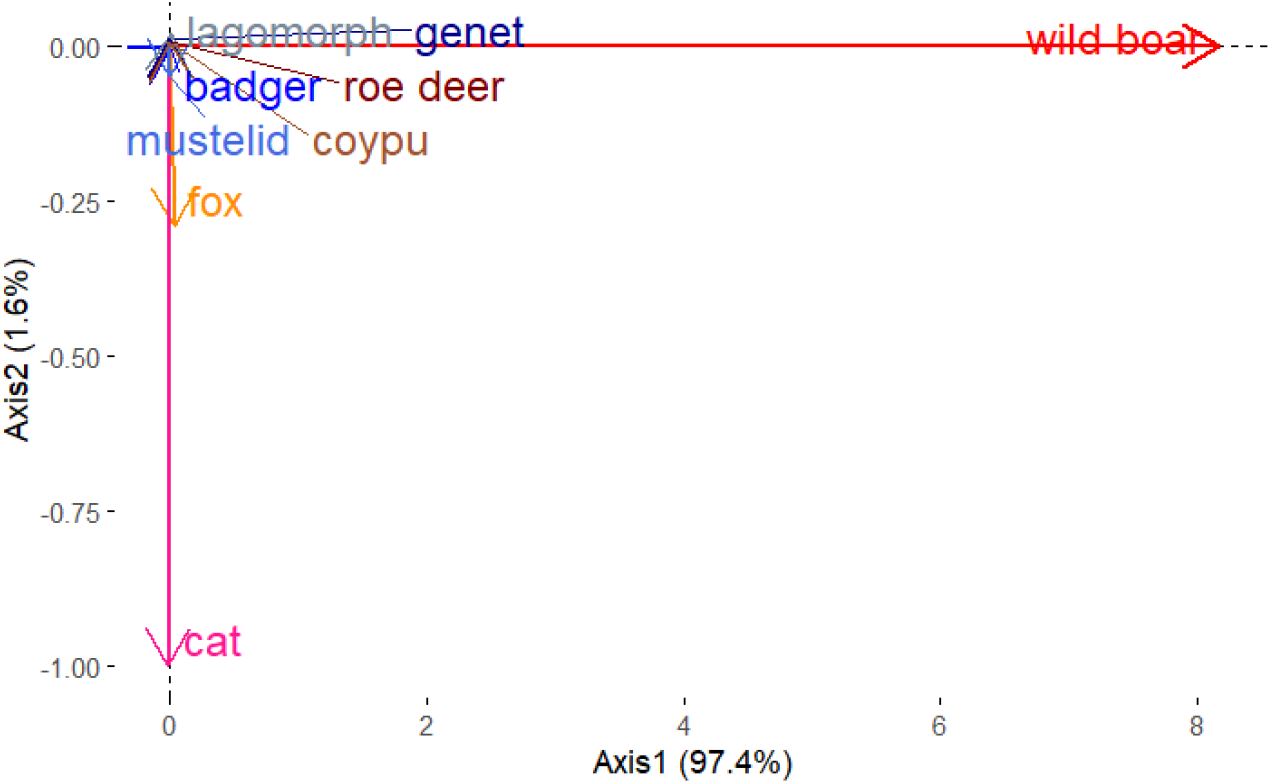
Biplot of principal component analysis on non-scaled centered data.

### No clear patterns of covariation with other mammal species

The first two axes of the PCA on scaled and centered data captured only a moderate proportion of the total variability of the data, as the first two axes explained 33.7% of the total inertia. Axis 1 contributed 18.6% and axis 2 contributed 15.1 % (Fig. 4). The species that contributed the most to the first axis were mustelids (38.3%) and genets (20.3%). Although roe deer contributed less to this axis, their contribution was opposed to that of all other species. The species that contributed the most to axis 2 were badgers (29.2%), lagomorphs (27.4%), and cats (21.9%). Wild boar exhibited minimal contribution to either of these axes (0.3% for axis 1, 1.9% for axis 2).

**Fig. 4.**
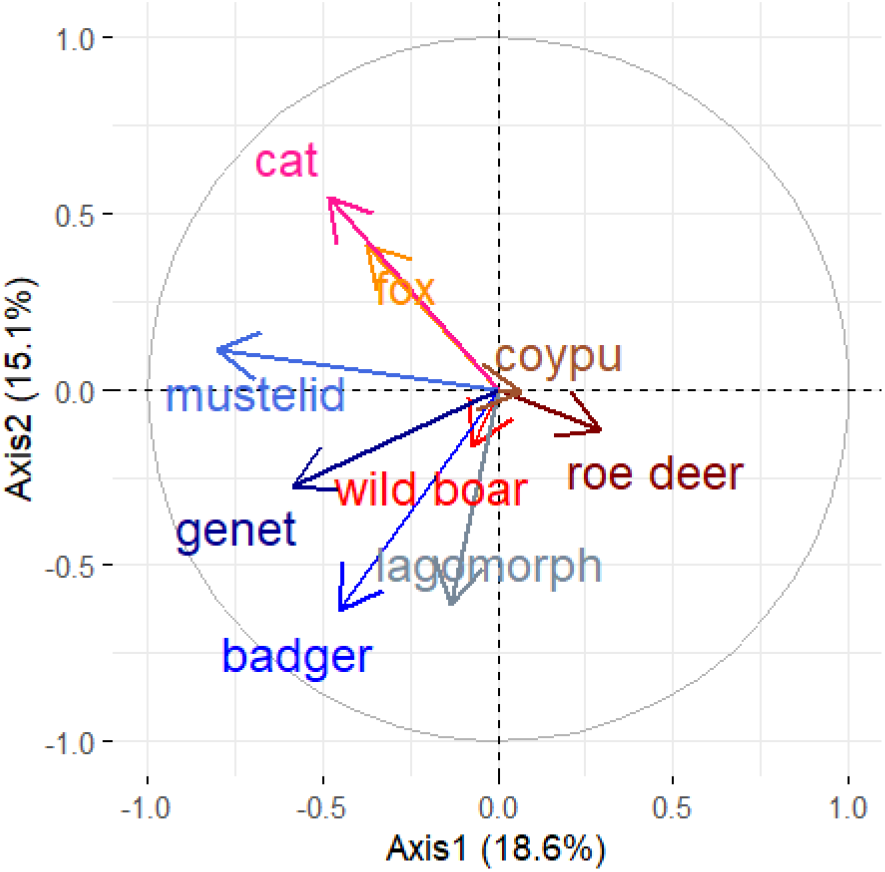
Biplot of the principal component analysis on scaled centered data.

### Spatial and temporal variations

The representation of CTs on the PCA scatter plot showed larger and less overlapping ellipses than the representation of months (Fig. 5). This suggests that spatial variations (i.e. differences between CTs) contributed more to the overall variability observed in the data set than temporal variation (i.e. differences within CTs). In particular, some CTs were very different from the others (Fig. 5a). The between-and within-group PCAs supported this interpretation.

**Fig. 5.**
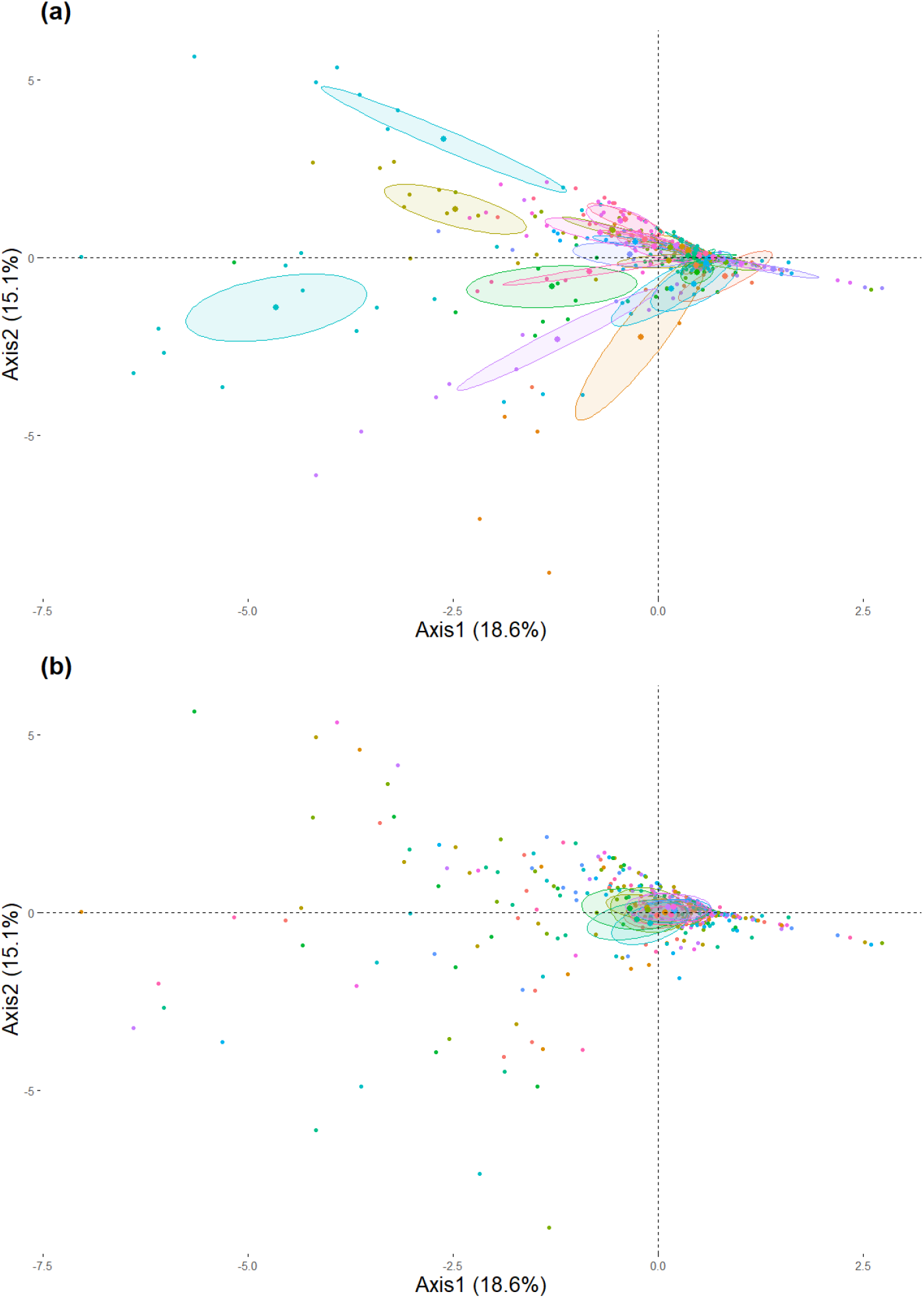
Scatterplot of the PCA on scaled centered data results on the first two axes. Each point represents the monthly weighted number of mammals for a CT, an ellipse represents the grouping of individuals from the same (a) CT (b) month, with a probability of 95%.

### Spatial variability : between-CT and within-month PCAs

The **between-CT analysis**, that focused on the differences between CTs (spatial), revealed that the CT variable explained a significant portion of the total variability in the dataset. The variability explained by the CTs amounted to 43.9% of the total variation, suggesting that the spatial setup or environmental conditions at different CTs locations significantly contribute to the observed variation in the data. Of the variability explained by the CT sites, respectively 30.9% and 22.2% were explained by axis 1 (driven by relative abundance of mustelids and genets) and axis 2 (driven by relative abundance of cats and foxes) of the between-CT PCA (Fig. 6). The scree plot shows a clear elbow point between axis 2 and axis 3, indicating that subsequent components contribute minimally to the data’s variance and can be disregarded when interpreting the results (Fig. 6a).

**Fig. 6.**
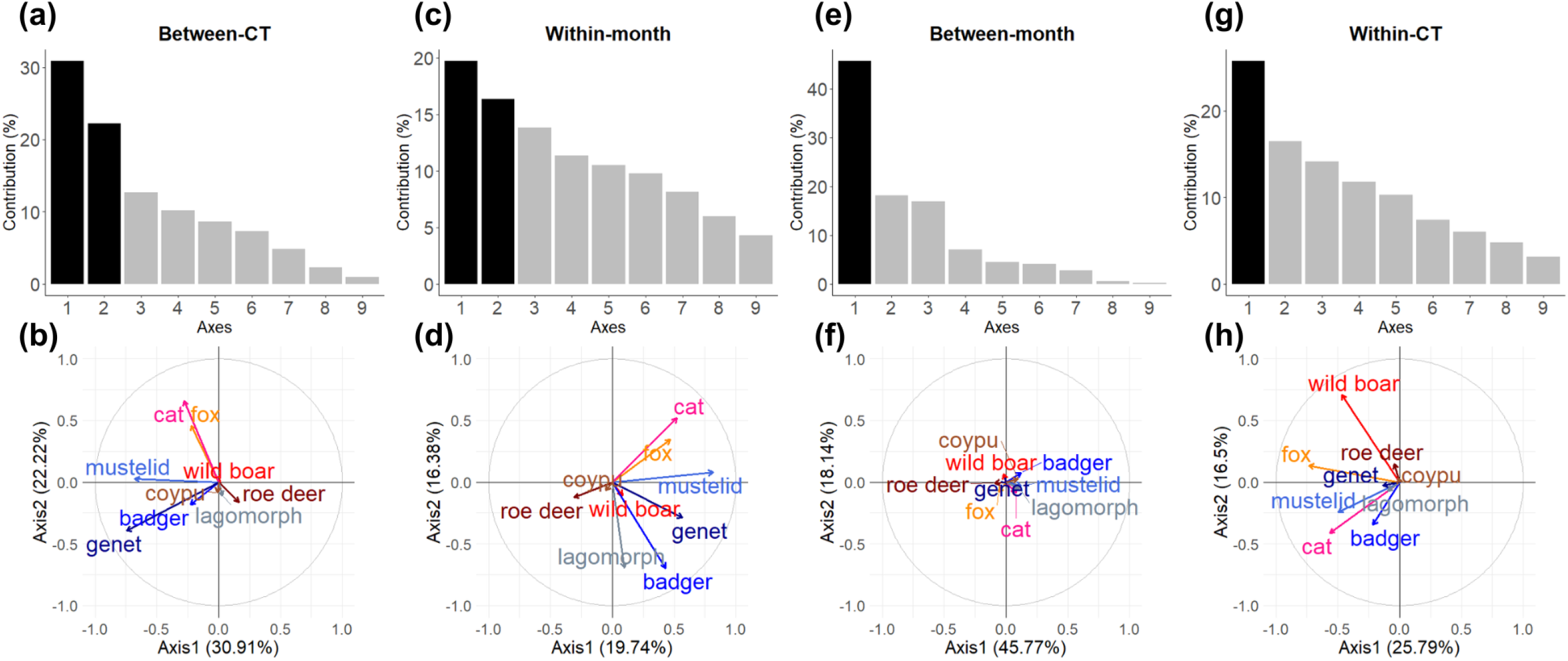
Results of the between-CT, within-CT, between-month and within-month principal component analyses. The scree plots, representing the percentage of contribution of the axes to the total inertia are shown in panels (a,c, e, g). The black bars represent the first principal components that explain the majority of the variance. Scatterplots of the results of the PCAs on the first two axes are shown in panels (b,d, f, h).

The **within-month analysis**, that removed the effect of months, showed that the variability observed in the dataset was not explained by consistent monthly variations. Indeed, the within-month variability accounted for 98.3% of the total variability, showing clearly that factors other than months, such as spatial differences, were the most important. In addition, the contribution of species to axes 1 and 2 somewhat matched the one observed in the between-CT PCA, with cat and foxes opposed to genets and mustelids on axis 1 (which accounts for 19.7% of the variance explained by this analysis), and roe deer were opposed to other species on axis 2 (which accounts for 16.4% of the variance explained by this analysis).

### Temporal variability : between-month and within-CT PCAs

As strongly hinted at by the results of the within-month PCA (above), the **between-month** analysis confirmed that the “month” factor explained only 1.7% of the total variance. As this was negligible, there was no need to interpret further the contribution of species to the axes (but see Fig. 6 if interested).

The **within-CT analysis**, which focused on the variability observed at the level of each CT site, showed that 57.8% of the total variability was attributable to differences that were not linked to differences between CT sites. This highlights an important level of within-site heterogeneity that, given the way the dataset was built, reflects a variability between months that is not consistent across species (given that between-month PCA explained a negligible share of variability). The scree plot (Fig. 6) showed that the first axis of the within-CT analysis was the only one to consider. It accounted for 25.8% of the variability explained by this analysis. All species contributed to this axis in the same direction, revealing that it represented an axis of general abundance of the studied species at the CT, to which the most common species (wild boar, cat, fox) contributed slightly more than others. This likely reflects the fact that some months had fewer sightings than others at each CT.

## Discussion

Our study demonstrates the extensive presence of wild boars in the peri-urban and urban areas of Montpellier. More importantly, it offers novel insights into the covariation between wild boar abundance and that of other mammals in these environments. We actually did not identify any consistent covariation between wild boars and specific species, suggesting that the strong spatial variability in wild boar abundance is likely linked to factors that affect other species differently. Overall, these results provide useful information for debates about the pros and cons of bush clearing to manage wild boars in urban areas. They reveal that it is difficult to make generalizations about which other species could be impacted, and highlight the importance of conducting short pre-interventions local studies.

### Wild boars are abundant in Montpellier

The deployment of CTs was primarily focused on areas where the presence of wild boars was suspected, based on observations of recent signs of presence (e.g. spoors) or information from municipal authorities or local residents. We assumed that these locations, which were presumed to be areas of high wild boar traffic, were indeed areas that could possibly be targeted by authorities for bush clearing interventions.

As anticipated given the way we selected CT sites, wild boars were detected in almost all sites (92%). Despite the bias in site selection, and the fact that the sites were widely distributed across Montpellier and neighbouring municipalities, our study thus confirms the widespread presence of wild boars in this city. This is particularly true given that, of the four sites where no wild boars were recorded, only one was active throughout the entire study period (52 weeks), while the other three were active for less than 10 weeks, meaning we could thus have missed some wild boar passage. In addition, at many sites wild boar was by far the most common species observed. Overall, our observations are consistent with the increasing recognition that wild boars can successfully colonize, and even thrived in, urban areas (Amendolia et al., 2019; Cahill et al., 2012; Licoppe et al., 2013; Stillfried et al., 2017b). A number of life-history traits seems to underlie this adaptability : wild boars have a remarkable dietary (Ballari and Barrios-García, 2014; Schley and Roper, 2003) and behavioural (Johann et al., 2020; Stillfried et al., 2017a) flexibility. For instance, in town, they can feed on both natural vegetation and human food waste (Ballari and Barrios-García, 2014; Castillo-Contreras et al., 2018), forage at night when there are less people and cars in the streets (Podgórski et al., 2013), and rest during the day in very small patches of vegetation and small ditches (pers. obs.; Fradin and Chamaillé-Jammes, 2023). In addition, life in the city also brings benefits: on top of human food waste that they access by taking down and opening bins, wild boars are sometimes directly fed by people and can easily access food provided to cats and pigeons (pers. obs., and social surveys associated with this study and in progress). Finally, as no hunting occurs, wild boars are not directly threatened by people, which might bring some physiological benefits through the loss of fear. It also facilitates their habituation to human presence (and sometimes dogs, pers. obs, and social surveys associated with this study), which is evident in some (but not all) wild boars encountered in the city (pers. obs.; Cahill et al., 2012; Stillfried et al., 2017a). Although we do not yet understand the relative contribution of these mechanisms in explaining the success of wild boars in Montpellier, our study clearly demonstrates that this city is to be added to the growing list of towns that need to consider human - wild boar coexistence.

### Local wild boars abundance does not covary with that of other species

Although wild boars are generally abundant in Montpellier, they are not distributed uniformly within the city, and we observed large variations in abundance at local levels. Explaining these variations was not the focus of the study, rather it aimed at understanding whether these variations were matched in other species, i.e. understanding whether sites with relatively more wild boars also had relatively more other species than other sites. By addressing this issue, we fill an important knowledge gap, as this topic has been poorly addressed in the literature, in particular in urban areas and in the native range of wild boars (Barrios-Garcia and Ballari, 2012).

Overall, we found through the between-CT PCA that local wild boar abundance did not spatially covary with those of the other species studied here. This was somewhat unexpected as we would have thought that a number of species, which would include wild boars, would covary positively, as there was a relatively high environmental heterogeneity, for instance in vegetation cover or building density, across our CT sites. We expected that a number of species would for example favor sites with more cover and avoid those with higher building density. Clearly, however, variations in wild boar local abundance were unrelated to those of other species. The mammal community studied appeared to be mostly structured in space by variations in the distribution of small carnivores. Further studies will look at the drivers that explain the distribution of wild boars, and possibly of some other species. However, our results may suggest that the great flexibility of wild boars ‘decouples’ them from the environment more than other species (for instance compared to mustelids and the common genet) and possibly differently to other flexible species, such as cats and foxes, due to their diet. This is particularly relevant here as our CTs mostly recorded wild boars in active areas, where they are likely to exhibit lower environmental selectivity than at their resting sites.

Although the mammal community studied has clearly spatially structured over our area of interest, some significant temporal variability was also present. Our results (i.e. the between-month PCA) did not reveal a consistent, community-level, change across months. This outcome was expected given the interspecific variability in life history traits and trophic strategies among the studied species. Rather, results highlighted, through the within-CT PCA, that, for some species, the number of animals seen by any CTs could vary quite importantly between months. This could be caused by, for instance, seasonal disturbances (increased human presence in parks during spring and summer, drive hunts in autumns and winters in some peri-urban areas etc.) and reproductive schedules which could both cause redistribution of animals in the landscape and thus substantial variations in the number of sightings obtained monthly at each CT site.

We cannot rule out however that some of this variation could have occurred at a much more local scale, with animals slightly changing their commuting habits, for instance by using another path than the one passing in front of the CT. The small field-of-view of CTs make the collection of data sensitive to small-scale choices made by animals (Burton et al., 2015; Hofmeester et al., 2019), and ideally high-density CT grids should be used to reduce this effect (Rovero et al., 2013). For a given number of CT, it would however force reducing the number of sites studied. Here, we favored covering a large number of sites distributed throughout Montpellier in order to study the spatial structuring of the community at this scale.

### Management implications

Our study directly arose from a management-oriented question we were asked in the context of a larger program on the ecology of wild boars in urban areas. The question was: “Is bush clearing in areas frequently by wild boars likely to affect other species, and if so, which ones?”. Our results answer this question: locally, bush clearing intervention could affect a number of other species, in particular small carnivores such as the common genet which is a legally-protected species in France. However, because of the lack of spatial covariation between wild boars and other species, we cannot reliably predict which species would be likely affected. It could very well be genets in one area, mustelids in another, and neither species in a third place. This is a disappointing answer from a management perspective, as it suggests that bush clearing interventions would be conducted blind to their effects, or that a local biodiversity evaluation would be needed prior to any intervention. Given that this latter preferred option would be more costly, we recommend concentrating interventions on sites where the presence of wild boars has been problematic, rather than applying them generally under the assumption that wild boar presence is inherently detrimental. Note that our results of a lack of spatial covariation between wild boars and other species does not support the idea that wild boars are a dominant negative driver of the abundance of the species studied here. Therefore, a spatially-targeted intervention approach appears to be the best way to address wild boar management issues in the context of hard-to-predict impacts of these interventions on biodiversity.

## Acknowledgements

Lucie Favier contributed for a short period to the fieldwork and to some preliminary analyses. We acknowledge the support of a number of people that helped J. Dezeure with the authorization to deploy camera-traps. In particular, we thank Mr Thaler, Mr Gartner, la Maison de la Nature of Lattes, and the municipalities of Montpellier, Castelnau-le-Lez, Saint-Aunès, Montferrier-Sur-Lez, Grabels, Juvignac, Lavérune, Saint-Jean-de-Vedas and Villeneuves-les-Maguelones for their collaboration.

## Author’s contribution

S. C.-J., R. M, S. S. and E. B. designed the study. J. D. conducted the data collection and managed the data. L. B. analyzed the data, under the supervision of S. S., E. B. and S. C.-J. L. B. wrote the first draft of the manuscript, which was commented and revised until all authors approved the final manuscript.

## Data availability

Data are available upon request to the authors for replicating the study, and discussions about collaborative work will be welcomed.

